# Induction and long-term maintenance of hindbrain-like neural stem cells in xeno- and basic fibroblast growth factor-free conditions

**DOI:** 10.1101/2025.03.03.640169

**Authors:** Ziadoon Al-Akashi, Denise Zujur, Nicholas Boyd-Gibbins, Nathalie Eileen Wiguna, Masato Nakagawa, Tetsuhiro Kikuchi, Asuka Morizane, Jun Takahashi, Makoto Ikeya

## Abstract

Neurons exhibit region-specific identities corresponding to functional distinctions across different brain areas. Region-restricted neural stem cells (NSCs) have previously been generated from pluripotent stem cells; however, maintaining their regional identity over extended passages remains challenging. Here, we report the generation of hindbrain-like induced NSCs (Hb-LiNSCs) with upregulated hindbrain-specific markers and downregulated forebrain, midbrain, and spinal cord markers under xeno- and basic fibroblast growth factor-free conditions using three chemicals—CHIR99021 (at a high concentration), a potent activator of the Wnt pathway, A-83-01, a potent inhibitor of the TGF-β/Activin/Nodal pathway, and LDN193189, a potent inhibitor of the bone morphogenetic protein pathway. Hb-LiNSCs maintained their chromosomal integrity, multipotency, and differentiation capacity even after long-term culture for more than 60 weeks. This innovative approach enhances our understanding of neurodevelopmental and neurodegenerative processes in the hindbrain region and paves the way for developing targeted cell-based therapy as well as disease modeling for drug discovery.

## INTRODUCTION

Neural stem cells (NSCs) are multipotent cells that arise during development and reside along the neural tube. These cells proliferate and differentiate into the various cell types that form the central nervous system regions during embryological development. NSCs from different regions of the neural tube express specific regional markers^1^. Since the first report on neural stem cells in the dentate gyrus of adult rat brains was published over five decades ago, significant progress has been made in isolating NSCs from various brain regions^2^. Primary and induced (iNSCs) NSCs are invaluable for studying neurogenesis and modeling neurological disorders.

Cells isolated from the hindbrain of aborted human early embryos were reported to maintain their regional specification and neurogenic proliferative capacity even after long-term culture^3^. Although such an approach provides sustainable, healthy donor-derived hindbrain NSCs, isolating patient-derived hindbrain NSCs from fully developed humans is prone to a great risk of damaging neurons passing through the brainstem that innervate vital organs. Furthermore, the use of human embryos raises ethical concerns.

The use of human induced pluripotent stem cells (hiPSCs) could overcome the limitations associated with NSC isolation from adult patients and human embryos. hiPSCs provide a scalable source for virtually any cells in the human body, and the ability to obtain hiPSCs from multiple somatic tissue sources renders them useful for disease modeling and regenerative medicine^4^. The induction of NSCs from several clones of hiPSCs derived from different tissues has been evaluated in previous studies. Dual SMAD inhibition was used to induce mid/hindbrain type NSCs regardless of their somaticorigin^5^. Moreover, activation of WNT signaling was shown to control the rostral–caudal neural axis in a dose-dependent manner using a microfluidic culture device controlling a highly potent and selective GSK inhibitor (CHIR99021), and the development of neural tube expressing the forebrain, midbrain, and hindbrain markers could be modeled using this system depending on the CHIR99021 concentration^6^.

In this study, we aimed to generate iNSCs from hiPSCs having regional specification restricted to the hindbrain (hindbrain-like iNSCs [Hb-LiNSCs]) by upregulating hindbrain gene expression and downregulating forebrain and spinal gene expression under minimal growth factor conditions. To achieve this, we performed a mini-screening and identified high concentration of CHIR99021, along with LDN193189, an inhibitor of bone morphogenetic protein (BMP) type I receptors ALK2/ALK3, and A-83-01, a potent inhibitor of the TGF-β/Activin/Nodal pathway that inhibits ALK5, ALK4, and ALK7, without the use of basic fibroblast growth factor (bFGF) nor serum. Using these minimum induction conditions, we could maintain Hb-LiNSCs for up to 60 weeks in cell culture. We characterized these Hb-LiNSCs during the early (5–20 weeks), mid (25–40 weeks), and late (45–60 weeks) passages and confirmed their long-term maintenance in culture.

## RESULTS

### Induction of Hb-LiNSCs

Hb-LiNSCs were induced from hiPSCs by inhibiting mesodermal and endodermal pathways through the inhibition of the TGF-β/Activin/Nodal pathway and BMPs using A-83-01 and LDN193189, respectively, and adding posterior characteristics by activating Wnt signaling using CHIR99021 (this condition is hereafter referred to as “ACL”). iPSCs were plated on hLaminin-511E8 fragment-coated culture wells in the same medium used for maintaining iPSCs but without bFGF and with ACL, adding Y-27632 on the first day (Figure 1A). During the first week of culture, multiple independent dome-shaped colonies, which were distinct from the typical confluent (flat) iPSC colonies, were formed. These dome-shaped colonies could spontaneously change their morphology, forming neurite-like projections, when ACL was removed from the culture medium (Figure 1B). The morphology of dome-shaped colonies was maintained after cell passage and was similar in the three different iPSCs lines tested (Supplementary Figure 1A).

**Figure 1.**
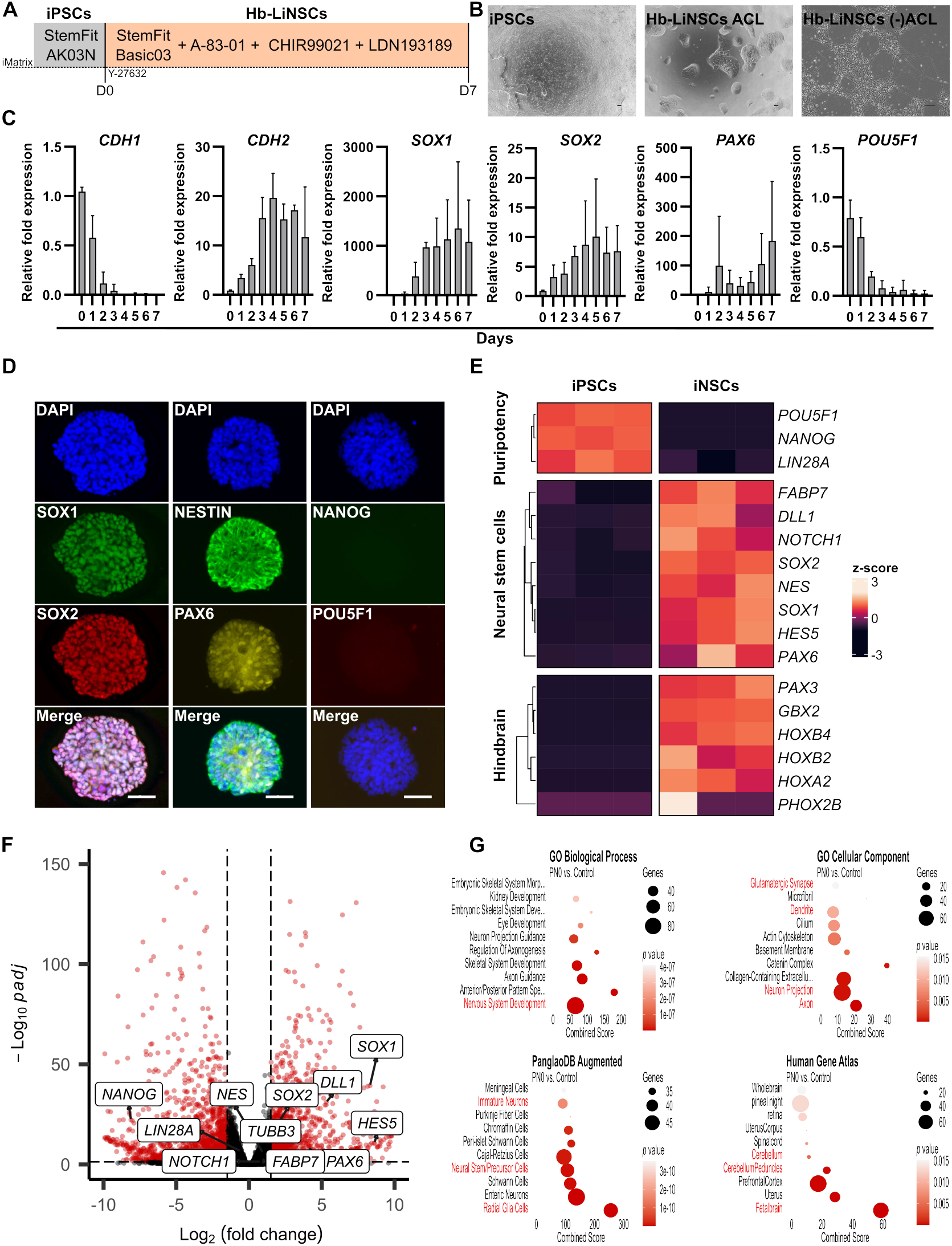
Induction of hindbrain-like neural stem cells. (A) Schematic of the method used to induce human induced pluripotent stem cells (iPSCs) into hindbrain-like neural stem cells (Hb-LiNSCs). (B) Phase contrast images of iPSC and Hb-LiNSCs colonies, and differentiated neurons from left to right. Scale bar, 100 µm. (C) E-cadherin (*CDH1*), N-cadherin (*CDH2*), *SOX1, SOX2, PAX6*, and *POU5F1* gene expression in iNSCs during the induction phase relative to that on day 0 (iPS cells). Bars indicate means; error bars as standard deviation, *n* = 3. (D) Immunocytochemical staining for SOX1, SOX2, NESTIN, PAX6, NANOG, POU5F1, and that of nuclei with DAPI in Hb-LiNSCs at day 7 after induction of iPSCs. Scale bar, 50 µm. (E) Heatmap of selected genes representing pluripotency, neural stem cells, differentiated neurons, and canonical markers for the hindbrain region, with hierarchical clustering of genes and samples. Three samples of 1231A3 iPS (as control) and 1231A3 Hb-LiNSCs at PN0 day 7 were used. Normalized gene expression data are represented by the color intensity of the row z-scores. (F) Log foldchange values (x-axis) are plotted against significance (negative log_10_ of *p*-adjusted value, y-axis) for PN0 day 7 Hb-LiNSC samples (three samples) vs. 1231A3 iPS control (three samples). Red color dots indicate significantly differentially expressed genes. Vertical dashed lines indicate the threshold of 1.5 log foldchange, and the horizontal dashed lines indicates the significance threshold (*p* = 0.05). (G) Gene enrichment analysis of the significantly (*p* < 0.05) upregulated (log_2_ foldchange ≥1.5) genes in the PN0 Day 7 Hb-LiNSCs vs. iPSC control comparison for selected datasets. Dot size indicates the number of genes overlapping with the dataset, color intensity indicates the significance (top 10 terms ordered by *p*-values), and the x-axis indicates the combined score calculated using Enrichr. Relevant terms are highlighted in red.

Within 3 days, the cells exhibited marked downregulation of E-cadherin (*CDH1*), which is primarily expressed in epithelial cells, and upregulation of N-cadherin (*CDH2*), which is characteristic of neurons. Additionally, the cells gradually upregulated the expression of NSC markers, namely *SOX1, SOX2*, and *PAX6*, and downregulated the expression of the pluripotency marker *POU5F1* (Figure 1C). After removing any one or two of the ACL small molecules, the colonies were still formed, and the cells survived through the first week (Supplementary Figure 1B). Even though the ectodermal markers were upregulated under these conditions, the mesodermal and endodermal markers were also upregulated. Therefore, the use of all three ACL molecules was deemed essential (Supplementary Figure 1C). The gene expression was confirmed to be consistent in iNSC colonies on day 7 of induction using immunocytochemical staining. The expression of NSC markers, but not of pluripotency markers, was detected (Figure 1D).

To further investigate the ACL induction condition, we utilized bulk RNA sequencing (RNA-seq) to examine the gene expression in three clones of the 1231A3-derived iNSCs for samples from day 7 after induction (PN0 D7). The pluripotency markers were downregulated but the known NSC and hindbrain regional markers were upregulated compared with that in the iPSC samples (Figure 1E, F). The pathway enrichment analysis of significantly upregulated genes using public datasets revealed significant enrichment of terms relevant to the nervous system development, such as “Nervous System Development” among the gene ontology (GO) biological process terms and “Glutamatergic,” “Dendrite,” “GABA-ergic,” “Axon,” and “Neuron Projection” among the cellular component datasets, highlighting the formation of progenitors with multiple neuronal subtypes along with the enrichment of “Immature Neurons,” “Neural Stem/Precursor Cells,” and “Radial Glia Cells” in the PanglaoDB Augmented, and “Cerebellum,” “Cerebellum Peduncles,” and “Fetalbrain” in the Human Gene Atlas datasets, indicating the immature stem/precursor with caudal regional identity of these neurons (Figure 1G). These results indicated the potential induction of hiPSCs into a regionally restricted population of neuronal progenitors, especially those exhibiting hindbrain NSC characteristics.

### Long-term maintenance of Hb-LiNSCs

To confirm whether Hb-LiNSCs were multipotent progenitor/stem cells, we sought to maintain them in long-term (up to 60 weeks) continuous cell culture, passaged and expanded every week. Every five passages, we cryopreserved subsets of cells and collected cell lysate for RNA analysis and repeated this process until week 60 (Figure 2A). This allowed us to have expandable cell stocks that could be thawed and used for further experiments and to obtain RNA samples at different time points during the long-term maintenance. To confirm that the chromosomal integrity was intact after long-term maintenance, we performed karyotyping and identified no changes even after 53 weeks (benign inversion of chromosome 9 that was observed was preexisting in the 1231A3 iPSC line) (Figure 2B, Supplementary figure 2A). To check whether the expression of NSC markers was preserved after the long culture, we performed immunocytochemistry at PN60 and confirmed that the cells uniformly expressed SOX1, SOX2, PAX6, and NESTIN, but not NANOG (Figure 2C). We then subdivided the samples into three groups of early-, mid-, and late-passage number Hb-LiNSCs based on passage number (time), and performed RNA sequencing with samples differentiated from early (PN5_diff) and late (PN60_diff) by removing the ACL small molecules and gradually changing the culture medium to neural differentiation medium (NDM). We plotted the results in a heatmap to assess the known genes for pluripotency, NSCs, and differentiated neurons, along with the known markers for the forebrain, midbrain, hindbrain, and spinal cord. Notably, the samples in the early group showed higher expression of the spinal cord and hindbrain markers. Some of these genes, such as *HOXB7* and *HOXB6*, showed a similar pattern of expression even after differentiation (Figure 2D). Despite these differences in expression, Hb-LiNSCs from early, mid, and late samples clustered together and showed slight variance in the principal component analysis (PCA) of the RNA-seq data (Figure 2E). This indicated that the long-term maintenance had a minimum impact on the gene expression profiles. To assess the impact of long-term maintenance on cellular identity, we analyzed the datasets used for Figure 1G, but only used the genes from the late-passage samples (PN45, PN50, PN55, and PN60) that were significantly upregulated compared with their expression in iPSCs. Similar processes and cell identity terms were enriched compared with those in the induction (in Figure 1F) or early samples (Supplementary Figure 2B). This consistency suggested that the neural stem/precursor state, with fetal hindbrain features, was maintained over time (Figure 2F). Further examination of gene expression patterns in Hb-LiNSCs from passage 5 to passage 60 revealed a high degree of correlation among the expressed genes (Figure 2G). This finding prompted us to further investigate the individual genes exhibiting differential expression across early, mid, and late passage Hb-LiNSCs.

**Figure 2.**
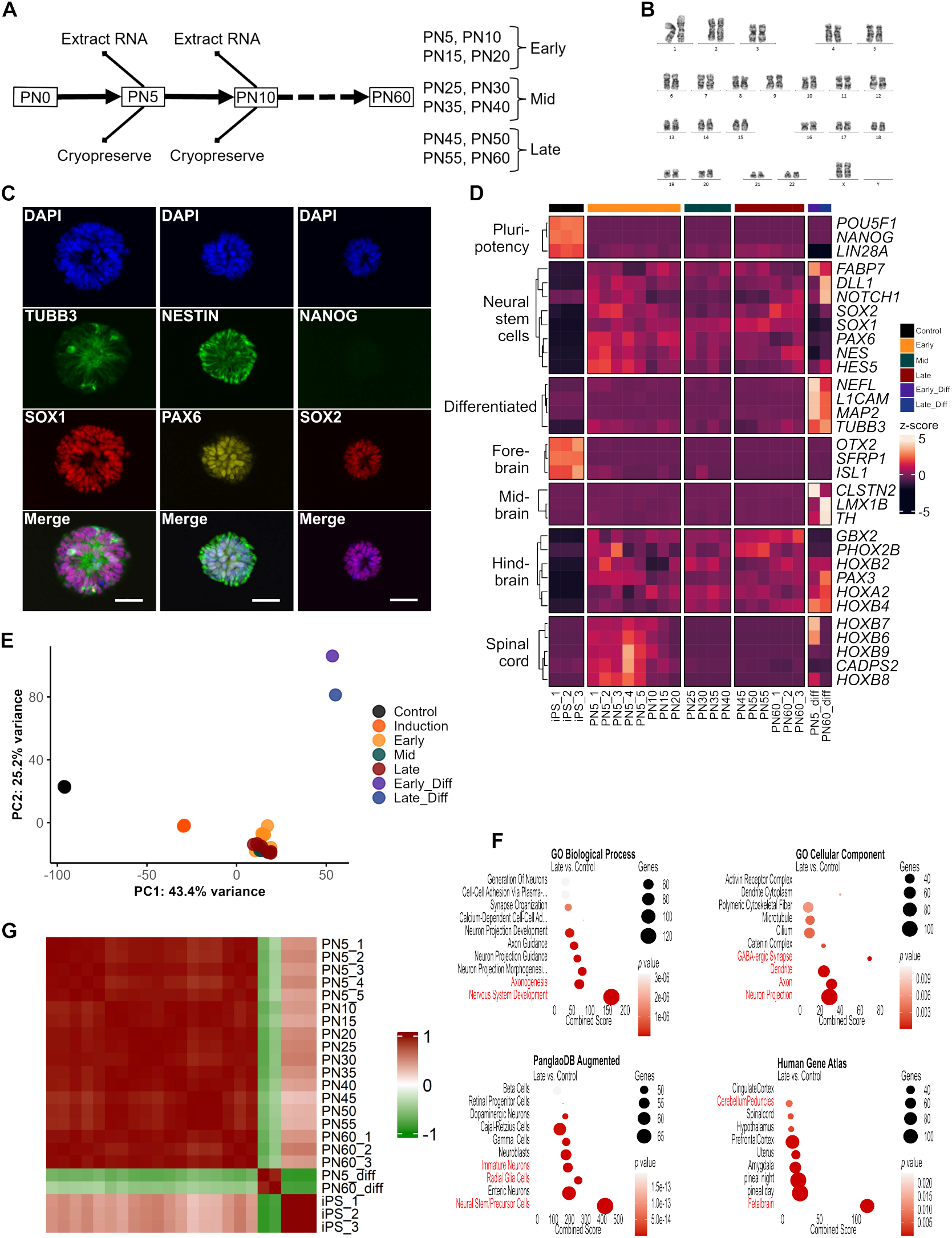
Long-term maintenance of hindbrain-like neural stem cells. (A) Schematic of the maintenance plan (on the left). Hindbrain-like neural stem cells (Hb-LiNSCs) were passaged every week, and thereafter, at every fifth passage, one clone of cells was cryopreserved, one was used to extract bulk RNA, and one was used for maintaining the culture. The grouping into early, mid, and late samples according to the passage number is shown on the right. (B) Karyotyping of Hb-LiNSCs at PN53 derived from the 1231A3 human iPSC line. (C) Immunocytochemical staining of Hb-LiNSCs for TUBB3, SOX1, NESTIN, PAX6, NANOG, and SOX2, and that of nuclei with DAPI at PN60 (60 weeks) after induction of iPSCs. Scale bar, 50 µm. (D) Heatmap of selected genes representing pluripotency, NSC, neural differentiation, and canonical markers for different brain regions, with hierarchical clustering of genes. The colored bar on the top indicates the samples. Samples PN5_1 to PN5_3 and PN60_1 to PN60_3 were derived from the 1231A3 iPSC line. Samples PN5_4 and PN5_5 were derived from HLAKO and SgT5 iPSC lines, respectively. “_diff” stands for differentiated neurons. Normalized gene expression data are represented by the color intensity of the row z-scores. (E) Principal component analysis (PCA) plots of the first and second (PC1 and PC2) components for the iPSC control are indicated in red, and Hb-LiNSCs and their differentiated cells at early-, mid-, and late-passage numbers are indicated in green, purple, blue, respectively. (F) Gene enrichment analysis of the significantly (*p* < 0.05) upregulated (log_2_ foldchange ≥1.5) genes in the late PN group vs. iPSC control comparison for selected datasets. Dot size indicates the number of genes overlapping with the dataset, color intensity indicates the significance (top 10 terms ordered by *p*-values), and the x-axis indicates the combined score calculated using Enrichr. Relevant terms are highlighted in red. (G) Pairwise correlation heatmap showing the relationships among samples. The color intensity in the heatmap indicates the correlation, ranging from 1 (positive correlation) in red through 0 (no correlation) in white to −1 (negative correlation) in green.

### Transcriptional differences between early and late passage Hb-LiNSCs

By comparing samples from early-, mid-, and late-passage numbers, we observed a gradual shift toward stabilization (Figure 2D). In the early vs. late comparison, we observed 1225 significantly differentially expressed genes (SigDEGs), of which 408 were upregulated (≥1.5-fold change); in the early vs. mid-passage sample comparison, we identified 547 SigDEGs, of which 207 were upregulated; whereas in the late vs. mid-passage sample comparison, only 153 SigDEGs were identified, with only 8 being upregulated (Figure 3A, Supplementary Figure 2C). Similar gradual decrease in the numbers of upregulated genes was observed when we compared early vs. control, mid vs. control, and late vs. control samples (Figure 3B). Although most of the significantly upregulated genes (1835 genes) were common between early, mid, and late samples, we could still observe the highest number of unique genes (225 genes) in the early vs. control comparison. This indicated that the early sample was more distinct from the mid and late samples. Therefore, we looked at the expression of several canonical genes for the embryological development of germ layers and regional marker genes. A similar level of downregulation of genes related to the mesoderm, endoderm, and pluripotency was noted. *HOXB9* was the gene with the highest expression in our list in the early vs. control comparison, whereas its expression was lost along with that of *HOXB8* and *HOXB7* in the late vs. control comparison (Figure 3C, D). *HOX 1–4* genes correspond to the hindbrain regions, whereas *HOX 5–13* correspond to the spinal cord. Furthermore, we examined the expression of *GBX2*, which is a transcription factor in more rostral regions of the hindbrain, and of *HOXB4*, which represents the more caudal regions of the hindbrain along with that of *SIX3*, which is a transcription factor regulating the forebrain, and of *HOXB9* as a marker for the spinal cord^7-9^. Significant differences were observed in the expression of the hindbrain markers between samples from early and late passages; however, both showed upregulation relative to that in RNA samples derived from adult healthy human pons and cerebellum (Figure 3E). Although the expression of *SIX3* remained comparatively the same and lower than that in the human-derived sample, *HOXB9* showed a significant difference between early and late passages even though the expression was still markedly lower than that in the human RNA sample from the spinal cord. To further understand the significance of differences in the early samples, we performed a pathway enrichment analysis for the genes that were significantly upregulated and unique to only the early vs. control group comparison (the 225 genes shown in the Venn diagram in Figure 3B). These unique genes showed enrichment of “Anterior/Posterior Pattern Specification” in the GO biological process but less significant enrichment of the other terms, namely “Immature Neurons” in the PanglaoDB Augmented dataset and “Fetalbrain” in the Human Gene Atlas dataset (Figure 3F). Upon direct comparison of the groups against each other, we found the highest number of unique genes in the early vs. late group comparison. We looked at these genes and found that *HOXB6-9* was consistently among the top 15 SigDEGs (Supplementary Figure 2D-F). These findings may indicate that the changes occurring after long-term maintenance are less likely to be related to maturation but more likely to be region-restricted specifications.

**Figure 3.**
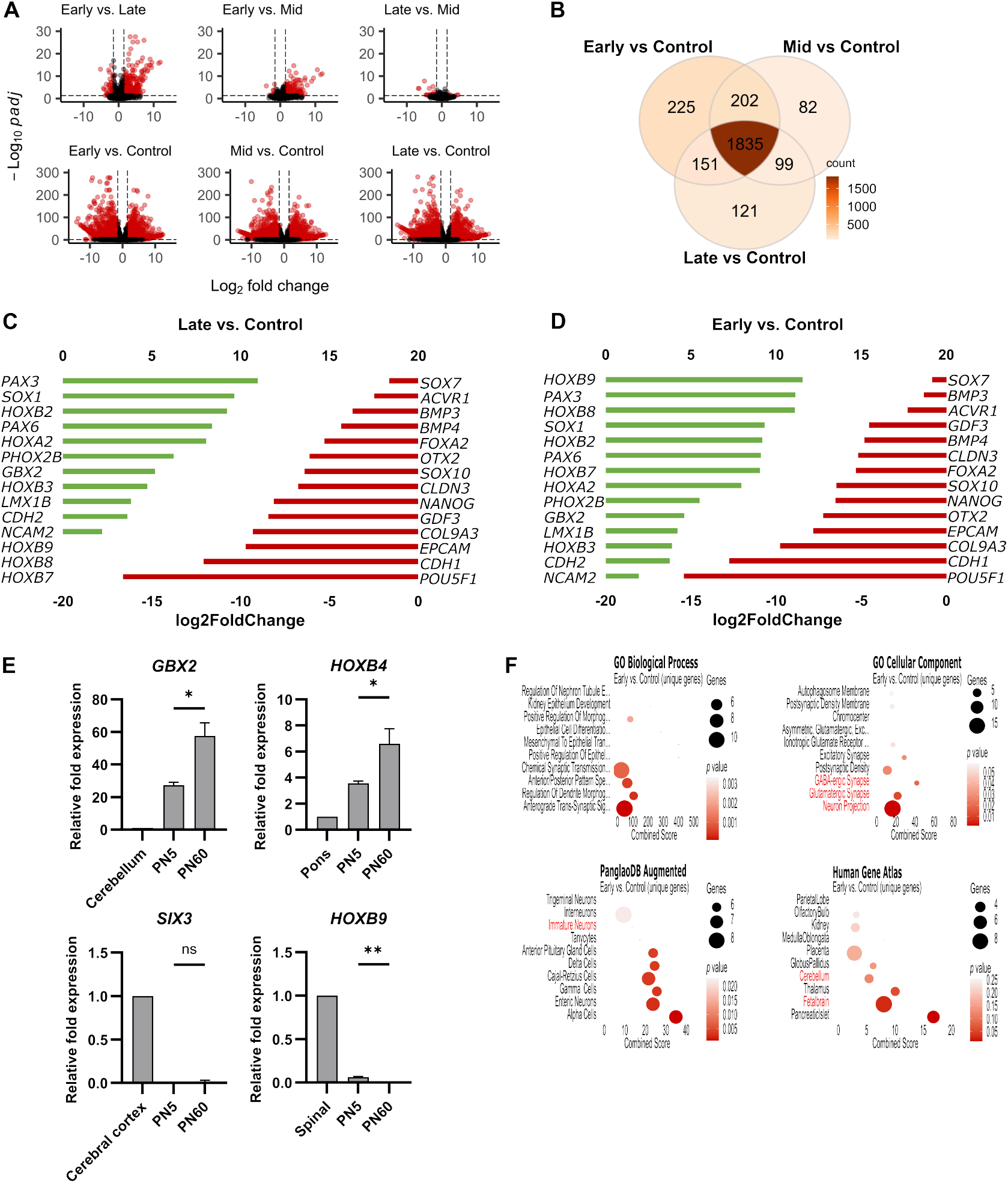
Transcriptional differences between early and late passage hindbrain-like neural stem cells. (A) Log foldchange values (x-axis) are plotted against significance (negative log_10_ of *p*-adjusted value, y-axis) for Hb-LiNSCs samples at one phase of maintenance vs. another (top panel) and for Hb-LiNSCs samples at different phases against the 1231A3 iPSC control (bottom panel). Red color dots indicate significantly differentially expressed genes (SigDEGs). Vertical dashed lines indicate the threshold of 1.5 log foldchange, and the horizontal dashed line indicates the significance threshold (*p* = 0.05). Number of SigDEGs is indicated on top of the plots. (B) Venn diagram of the significantly (*p* < 0.05) upregulated (log_2_ foldchange ≥1.5) genes in the Hb-LiNSC sample at different phases. Color intensity indicates the count of genes (darker color indicates a higher value). (C) Bar plot of the log foldchange of selected genes from the “Early vs. Control” bulk RNA-seq differential gene expression data. On the left, the list of genes with bars in green indicates upregulated expression (top x-axis). On the right, the list of genes with bars in red indicates downregulated expression (bottom x-axis). (D) Bar plot of the log foldchange of selected genes from the “Late vs. Control” bulk RNA-seq differential gene expression data. On the left, the list of genes with bars in green indicates upregulated expression (top x-axis). On the right, the list of genes with bars in red indicates downregulated expression (bottom x-axis). (E) Bar plot of RT-qPCR data for *GBX2, HOXB4, SIX3*, and *HOXB9* in PN5 and PN60 HbL-iNSCs samples (*n* = 3) relative to human-derived RNA samples from the cerebellum, pons, cerebral cortex, and spinal cord (*n* = 1). Unpaired *t*-test with Welch correction. n.s. ≥ 0.05, \**p* ≤ 0.05, \*\**p* ≤ 0.01. (F) Gene enrichment analysis of the 427 unique genes in the “Early vs. Control” comparison for selected datasets. Dot size indicates the number of genes overlapping with the dataset, color intensity indicates the significance (top 10 terms ordered by *p*-values), and the x-axis indicates the combined score calculated using Enrichr. Relevant terms are highlighted in red.

### scRNA-seq of early- and late-passage number Hb-LiNSCs reveals distinct cell populations

To investigate the differences between early- and late-passage Hb-LiNSCs at a single cell level, we obtained samples at different time points (PN5, PN21, PN41, and PN59) to represent the early, early-mid, late-mid, and late passages, respectively, and performed scRNA-seq. Different but comparative cell numbers were used in the analysis for each sample (Figure 4A). To determine the heterogeneity within samples, we used the Uniform Manifold Approximation and Projection (UMAP) together with the Leiden clustering algorithm and identified 16 distinct clusters (Figure 4B). We then labeled the cells with their corresponding samples and found that all the clusters were present in all the samples except for clusters 6, 9, 10, 12, and 13, which were represented almost strictly by the PN5 sample (Figure 4C). We examined the expression of canonical marker genes in samples for the forebrain, midbrain, and hindbrain regions of the neural tube along with markers for NSCs and differentiated neurons, such as serotonergic, GABAergic, and dopaminergic, and others, which were designated as “differentiated.” The heatmap showed that the genes present in the highest number of cells were those related to the NSCs and hindbrain region, whereas spinal cord markers were strictly present in the PN5 sample (Figure 4D). Clusters 0, 1, 2, 3, 4, and 7 were in proximity and revealed the highest percentages of cells expressing markers for both the hindbrain region and NSCs. Moreover, the expression of *HOXB9* was almost exclusively expressed in the cluster corresponding to the PN5 sample, whereas *HOXB4, HOXB2, SOX1, NES, SOX2*, and *HES5* were expressed in almost all the clusters (Figure 4E). Similarly, the heatmap for the expression of the top 10 differentially expressed genes across samples revealed that *HOXB9* was the top differentially expressed gene in cells from PN5, followed by *HOXB7* and *HOXB8* at the third and further position, respectively (Figure 4F). As indicated by the fold difference in expression for the top 25 genes in the PN5 vs. PN59 comparison, many of the *HOX* genes were upregulated (Supplementary Figure 3A). In the comparison of each sample with PN59, the score was highest for the genes differentially expressed in the PN5 vs. PN59 comparison than in the PN21 or PN41 vs. PN59 comparison, highlighting the major difference being between PN5 and other samples; this difference gradually faded with increasing passage number (Supplementary Figure 3B, C). These results allowed us to re-label the clusters with their indicative cell types, highlighted by the largest cluster belonging to the hindbrain stem cells (Figure 4G). An analysis of the distribution of cell types within the samples showed that cells related to the spinal cord were almost entirely restricted to the PN5 sample (Figure 4H). These results may indicate that, although there is a population of cells expressing both the spinal cord and hindbrain marker genes, this population transitioned over time with long-term maintenance. However, this shift did not affect the number of cells expressing the hindbrain NSC markers, which remained the highest in all the samples (Supplementary Figure 3D).

**Figure 4.**
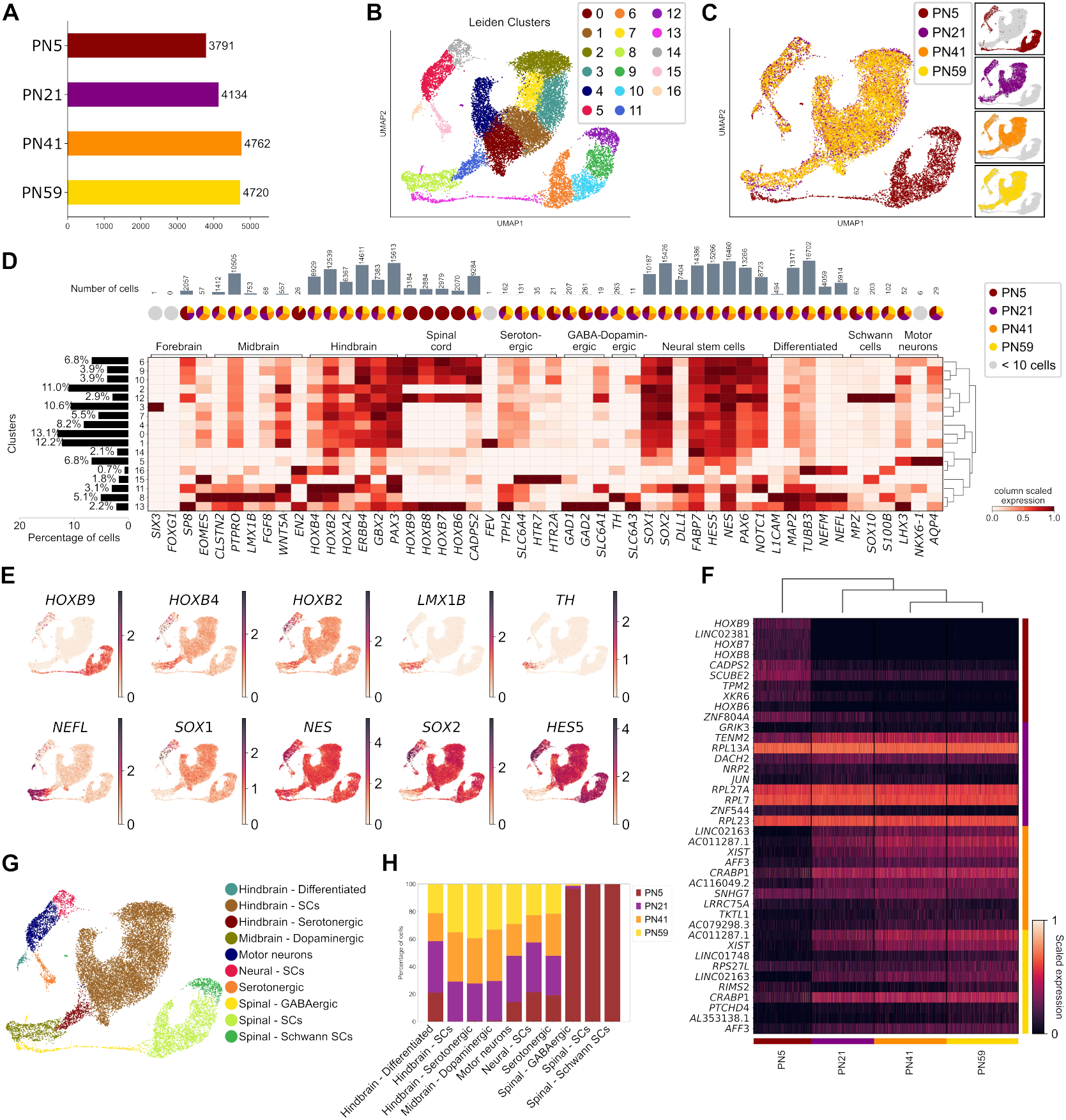
scRNA-seq of early- and late-passage number hindbrain-like neural stem cells reveals distinct cell populations. (A) Bar plot showing the number of cells in each sample ordered by passage number. (B) Uniform Manifold Approximation and Projection (UMAP) visualization of scRNA-seq data representing early (PN5), mid-early (PN21), mid-late (PN41), and late (PN59) hindbrain-like neural stem cell (Hb-LiNSC) samples from individual culture wells. Each dot on the plot corresponds to a single cell and is colored based on cluster assignments obtained via Leiden clustering (see methods for more details). (C) Left: UMAP plot representing cells color-coded by their respective samples. Right: Individual subplots highlighting each sample. (D) Heatmap showing the relative gene expression across clusters identified via Leiden for canonical markers associated with the brain regions, neural stem cells, and differentiated neurons. Rows are hierarchically clustered. On the left side, the bar plot shows the percentage of cells in each cluster. At the top, the bar plot indicates the number of cells expressing each gene, and the pie chart indicates the proportion of each sample expressing each gene; grey circles denote genes expressed in fewer than 10 cells. (E) Expression of selected genes across the cells on the same UMAP in (A). (F) Heatmap illustrates the expression levels of the top 10 differentially expressed genes across sample groups. Rows represent genes, and columns represent samples, with the color scale indicating standardized expression from low (darker shades) to high (lighter shades). Dendrograms are included for gene and sample clustering. (G) UMAP representation of the same dataset in (A), with each cell color-coded by the specific cell types, determined via the expression of canonical genes. (H) Stacked bar plot showing the distribution of cell types across samples. Each bar represents a cell type, with colored segments indicating the proportion of cells from different samples.

### Differentiation and functional network activity

Neurospheres are a cluster of cells cultured in a 3-dimensional (3D) manner floating in low-attachment round bottom plates, which are commonly used to assess primary or induced neural cells (Figure 5A). We used the neurosphere method following the removal of the ACL molecules and cultured the Hb-LiNSCs in NDM to differentiate them. We then performed attachment and differentiation analyses, electrophysiological recording, and in vivo implantation (Figure 5B). We seeded dissociated cells without any scaffolding material, and these cells formed sphere-like aggregates. These spheres were almost equal in size at initial seeding and exhibited significant growth after several weeks (Figure 5C, D). When Hb-LiNSC-derived neurospheres were attached to a flat surface coated with laminin, TUBB3-positive neurite-like projections were observed along with cells migrating outside the sphere edges (Figure 5E). Glial cells, such as astrocytes and oligodendrocytes, positive for GFAP and OLIG2, respectively, were observed. This differentiation capacity was maintained even after the long-term maintenance of Hb-LiNSCs.

**Figure 5.**
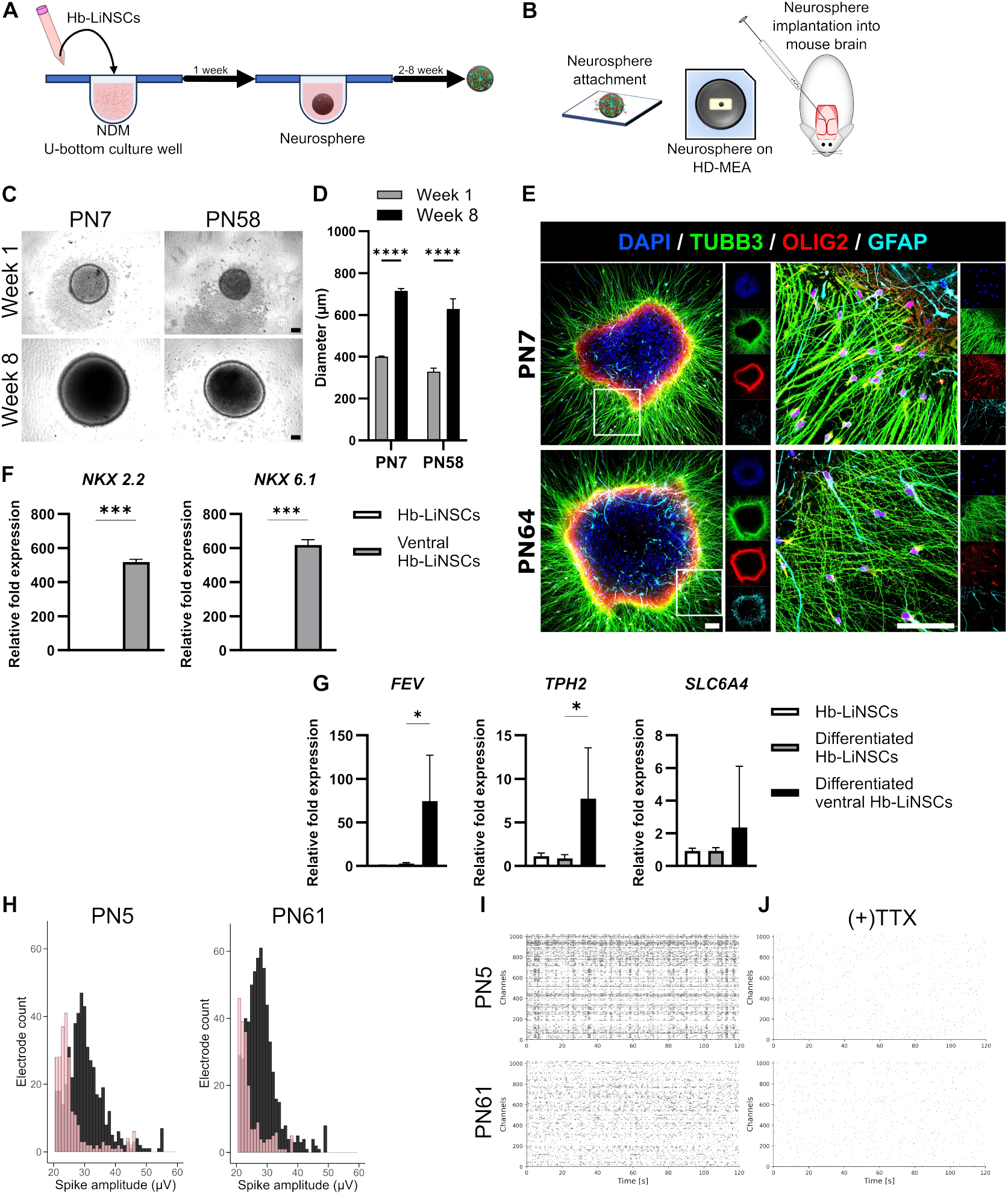
Differentiation and functional network activity. (A) Schematic representation of neurosphere formation. (B) The application of neurospheres in attachment to a flat surface, attachment to multielectrode arrays (MEAs), and in vivo transplantation. (C) Phase-contrast images of the neurospheres in suspension after one and 8 weeks of seeding. Scale bar, 100 µm. (D) Bar plot for the cross-sectional area of the neurospheres. (E) Immunocytochemical staining of the attached neurospheres for TUBB3, OLIG2, and GFAP, and for nuclei with DAPI. Scale bar, 100 µm. (F) RT-qPCR for gene expression of ventral hindbrain markers (*NKX 2*.*2* and *NKX 6*.*1*) in the ventralized hindbrain-like neural stem cells (Hb-LiNSCs) relative to that in the non-ventralized Hb-LiNSCs. (G) Gene expression for serotonergic markers (*FEV, TPH2*, and *SLC6A4*) for differentiated neurons relative to that in undifferentiated Hb-LiNSCs. Bars indicate means, with error bars showing the standard deviation, *n* = 3. (H) Histogram of the electrode counts for the detected spike amplitudes on the MEAs. The sample (PN5 left plot and PN61 right plot) before tetrodotoxin (TTX) treatment is shown in the dark, and that after TTX treatment is shown in pink. (I) Raster plot of the spike activity detected in the MEA overtime. Each dot represents an action potential (spike) for PN5-(top plot) and PN61-derived (bottom plot) samples. (J) Same samples as in (I)were recorded after adding TTX.

iNSCs can be further restricted to the ventral hindbrain region by treating the cells with sonic hedgehog (SHH) recombinant protein and FGF4, resulting in a population of serotonergic progenitor cells expressing *NXK 2*.*2* and *NKX 6*.*1* that can differentiate into serotonergic neurons expressing *FEV*, the serotonergic neurons transcription factor, *TPH2*, the tryptophan hydroxylate 2, and *SLC6A4*, a serotonin transporter, which are the key markers for serotonergic neurons found in the central nervous system ^10^. We treated the Hb-LiNSCs with smoothened agonist (SAG) along with FGF4 and obtained ventral Hb-LiNSCs expressing *NXK 2*.*2* and *NKX 6*.*1* at significantly higher levels than in the non-ventralized Hb-LiNSCs (Figure 5F). Upon differentiation of the ventral Hb-LiNSCs, the neurons significantly upregulated the expression of *FEV* and *TPH2*, but the difference in expression of *SLC6A4* was not significant when compared to that in non-ventralized Hb-LiNSCs (Figure 5G). This finding highlights the importance and the identity of Hb-LiNSCs as they can be ventralized and differentiated into highly specialized restricted type of neurons present in that region of the brain.

To evaluate the electrophysiological activity of Hb-LiNSC-derived neurospheres, we employed multielectrode arrays (MEAs). After one week of sphere formation, followed by 5–6 weeks of maturation and migration of the attached neurospheres to the electrodes, we detected bursts of action potentials (spikes) across multiple electrodes, with amplitudes exceeding 50 µV (Figure 5H). Although neurospheres derived from Hb-LiNSCs at later passages exhibited higher spike amplitudes across more electrodes, the firing rate was greater in those derived from Hb-LiNSCs at earlier passages (Supplementary Figure 4A). Additionally, repetitive firing was observed from some electrodes within milliseconds, followed by periods of rest and subsequent resumption of firing. Synchronous firing from multiple electrodes suggested the presence of network activity (Figure 5I). Upon adding tetrodotoxin (TTX), a neurotoxin that blocks sodium channels and thereby inhibits the conduction of action potential in neurons^11^, to the neurosphere culture, spike amplitudes were markedly reduced, and fewer electrodes exhibited activity compared to pre-TTX conditions (pink bars in Figure 5H). The frequency and network activity were also suppressed by TTX (Figure 5J, Supplementary Figure 4A). These findings indicated that long-term maintenance did not affect the differentiation potential, and Hb-LiNSCs could still generate functional neurons.

### In vivo transplantation

Finally, to investigate the integration and migration of Hb-LiNSCs in vivo, we transplanted one-week-old neurospheres derived from early- and late-passage Hb-LiNSCs into mice brains, interior to the cerebellum and into the pontine region (Figure 6A). After 8 weeks of transplantation, mice were sacrificed, and the integration and migration of the human neural cell adhesion molecule (hNCAM)-positive neural cells was detected using a human-specific NCAM antibody (Figure 6B). hNCAM-positive neural cells were detected in the spinocerebellar tract, cerebellum, corticospinal tract, and midbrain from samples derived from early- and late-passage Hb-LiNSCs (Figure 6C). The transplanted hNCAM-positive neural cells also showed TUBB3-, GFAP-, and OLIG2-positive cells (Figure 6D). These results indicated that the Hb-LiNSC-derived cells can differentiate and extend along with the host neurons, simulating normal physiological development and rendering useful as a tool for disease modeling and potential cell therapy.

**Figure 6.**
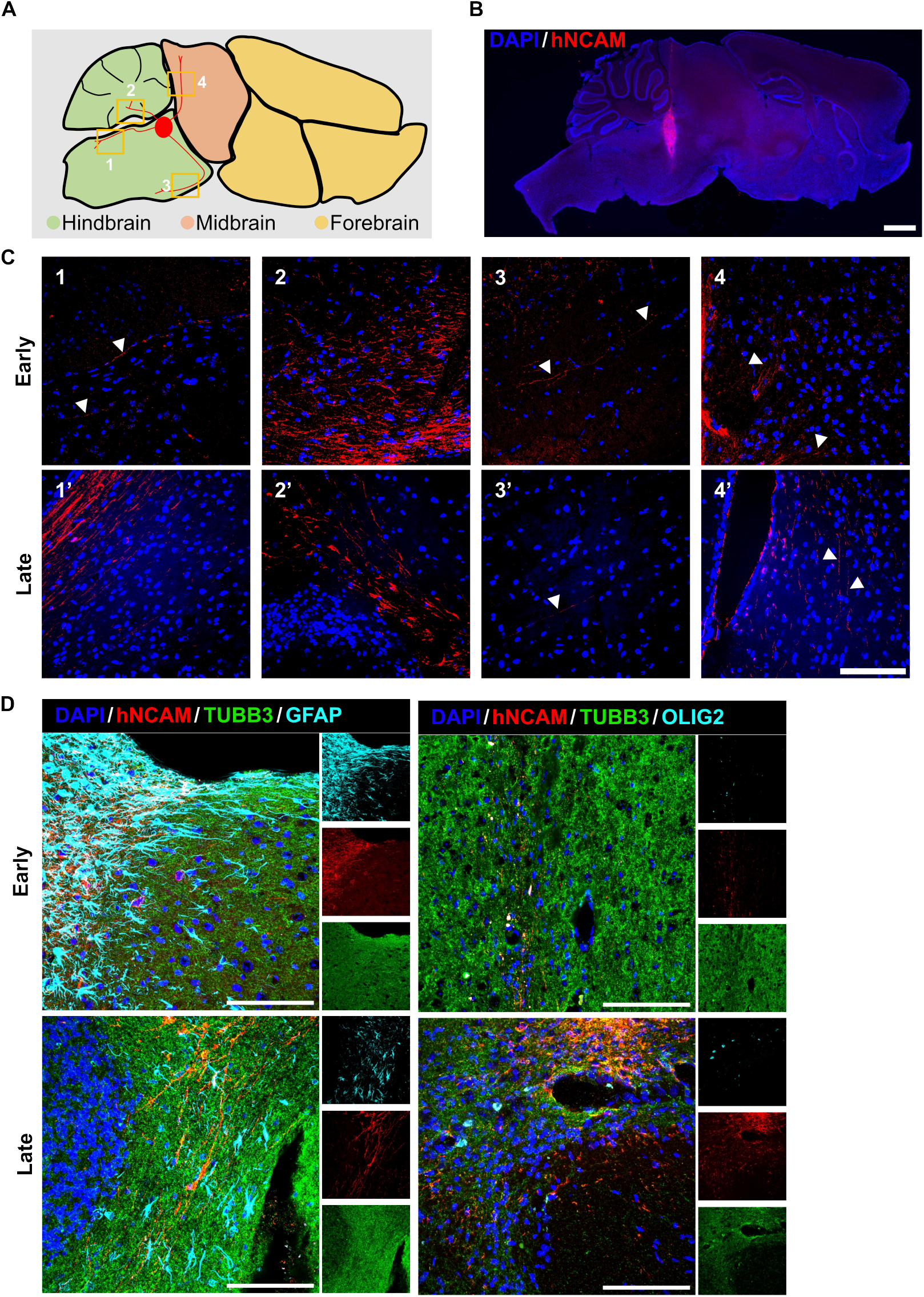
In vivo transplantation. (A) Schematic of the brain regions showing the transplantation site and the projection of neurite into the regions highlighted by orange squares and numbers. (B) Sagittal section of the mouse brain. (C) Tracking of human neural cell adhesion molecule (hNCAM)-positive cells along the spinocerebellar tract (1, 1′), cerebellum (2, 2′), corticospinal tract (3, 3′), and in the midbrain (4, 4′). (D) Differentiation of hNCAM-positive cells into TUBB3-, GFAP-, and OLIG2-positive cells in early- and late-passage samples.

## DISCUSSION

We successfully generated Hb-LiNSCs from hiPSCs under minimal induction conditions with small molecules without the addition of bFGF or serum. These Hb-LiNSCs could be maintained and passaged over a long term (we tested up to 60 weeks) without compromising on their chromosomal identity, genetic profile, differentiation potential, and functional activity. Our unique method adds valuable insights to the development of neural regional identity over long periods of stem cell maintenance, with cell passaging.

The regional specification is important in determining cell identity. The forebrain neurons are associated with higher cognitive functions, such as memory and learning; the hindbrain neurons regulate autonomic functions, and the spinal neurons are essential in relaying information from and to the brain. Moreover, pathologies arising from certain populations of neurons have dependencies on the region in question. This highlights the importance of using Hb-LiNSCs in disease modeling. One potential application of Hb-LiNSCs is in the modeling of certain types of malignancies referred to as diffuse interstitial pontine glioma (DIPG). In DIPG, a fatal tumor that affects children, the cells of origin are regionally restricted to the ventral pons^12^. Another potential application is in the modeling of disorders related to the serotonergic system. Serotonergic neurons are a highly specialized type of neurons that can utilize the neurotransmitter serotonin for their communication and cause disorders, such as major depressive disorder, autism spectrum disorder, Rett syndrome, and others. This potential can be expanded when Hb-LiNSCs from patient-derived iPSCs are established and then combined with MEAs to analyze their network activity, learning rate, and prediction accuracy by integrating neurons into a system that can analyze their spike (action potential) patterns data, such as the MEA-NAP, DishBrain, and BiœmuS^13-16^. Our Hb-LiNSCs have become a powerful research tool in this field, although there is a need for further improvement to leverage them for disease modeling and, potentially, drug screening.

We highlighted the differences between early- and late-passage Hb-LiNSCs, focusing on the gene expression evaluated using the sequencing data. We did bulk RNA sequencing of samples at different time points and compared data for those in the early induction phase and those maintained for the long term. Long-term maintenance did not affect maturation, and the terms related to immature/fetal neurons remained similarly enriched in both the early and late stages. scRNA-seq data for samples at different time points of maintenance revealed the largest number of cells (approximately 60%) expressing markers for the hindbrain region and coexpressing NSC markers. The second largest number of cells (approximately 14%) expressed spinal markers and coexpressed stem cell makers. Although the difference between early and late passages could not be fully understood, it is unlikely that maturation is involved; rather, based on the terms enriched for the differentially expressed genes, it is more likely that a higher regional realization occurs (Supplementary Figure 3E). Cells expressing spinal cord markers were only present in samples from early passages. This finding is not unique to our iNSCs; other researchers have shown that long-term maintenance results in higher expression of the forebrain and midbrain markers, probably due to CHIR99021 inhibiting the FGF targets, resulting in regression of the fate from the spinal cord^17^. The presence of the spinal cord markers in the early passage cells did not affect the other characteristics of NSCs with regard to gene expression, freeze/thaw, hindbrain regional markers, differentiation potential, and in vivo integration and migration. On the contrary, the electrophysiological activities were slightly different between the early and late passages. The number of electrodes in the MEAs detecting higher spike amplitudes was less for the neurons derived from early passage cells. In contrast, the frequency was higher for the samples derived from early passage cells when compared with the late passage cells. The underlying cause for this difference remains unclear, but it could be due to the early passage cell-derived neurons being regionally more heterogeneous, resulting in different network dynamics when compared with the late passage cell-derived neurons.

## RESOURCE AVAILABILITY

### Lead contact

Further information and requests for resources should be directed to the lead contact, Makoto Ikeya.

## Data and code availability

### Data

- Data reported in this paper can be shared by the lead contact upon request.

### Code

- The code for generating most of the figures can be found here: https://github.com/Ziadoon-Al-Akashi?tab=repositories

### Other

- Any additional information required to reanalyze the data reported in this paper is available from the lead contact upon request.

## ACKNOWLEDGMENTS

We would like to express our thanks to Fabian Raudzus, Takahiro Kitahara, and Edvinas Cerniauskas for scientific discussions, to Yoichi Kosodo for material provision, and to Yoshiko Inada, Kanichiro Nakano, Yukiko Nakagawa, Nakako Shimazu, and Misako Shimotsuji for administrative support. We thank the CiRA Foundation for single-cell RNA sequencing, the CiRA Common Equipment Management Office for providing the research instruments, such as qPCR and confocal microscope, and Marie Obien for technical support with MEA analyses. ZA would specifically like to express gratitude to the Otsuka Toshimi Scholarship Foundation. This work was supported by the iPS Cell Research Fund, grants-in-aid for scientific research from the Japan Society for the Promotion of Science (JSPS) (No. 16H05447), and Japan Agency for Medical Research and Development (AMED) under Grant Number JP15bm0104001 and 23bm1323001 to MI. This work was also supported by JST SPRING, Grant Number JPMJSP2110 to ZA.

## AUTHOR CONTRIBUTIONS

Conceptualization, ZA, MN, AM, and MI; methodology, ZA, DZ, NB, MN, TK, AM, and MI; investigation, ZA, DZ, NB, NEW, MN, TK, and AM; formal analysis, ZA, MN, and AM; writing – original draft, ZA. writing – review & editing, MN, TK, AM, JT, and MI; supervision: NB, MN, AM, JT, and MI; funding acquisition, ZA and MI

## SUPPLEMENTAL INFORMATION

**Document S1**. Figures S1–S4 and Table S1

## METHODS

### iPSC culture

We used the same iPSC lines and culture method indicated in our previous report ^18^. Briefly, iPSC lines 1231A3 (obtained from the Yamanaka laboratory), RPCiPS771-2 (SgT5-2) (Stemgent), and Ff-XT28s05-Abo_To (HLAKO) (CiRA Foundation), were cultured in iMatrix-511 silk (Nippi, 892021)-coated plastic 6-well plates in StemFit AK03N (Ajinomoto) under xeno-free conditions. The cells were passaged every week by dissociating them with Accutase^®^ (Sigma–Aldrich, A6964).

### Hb-LiNSC induction

After dissociation of iPSCs into suspension, 20,000 cells were plated in iMatrix-511-coated 12-well plastic plates in the same iPSC culture medium (AK03N) lacking bFGF, supplemented with 5 µM of A-83-01 (Tocris, 2939), 5 µM CHIR9902 (Axon, 1386), and 0.1 µM LDN193189 (MedChem Express, HY-12071) (referred to as “ACL”) along with 1 µM of Y-27632 (WAKO, 253-00513). The next day, the medium was completely changed with one without Y-27632. The AK03N medium without bFGF and with ACL was used for the maintenance of Hb-LiNSCs throughout the cell culture. Passage was carried out every week by dissociating the cells with Accutase, and Y-27632 was always added on the first day after plating.

### Neurosphere formation and Hb-LiNSC differentiation

Hb-LiNSCs were collected as dissociated cells, and approximately 10,000 cells were seeded in each well of 96-well U-bottom low-attachment plates in NDM consisting of Neurobasal™ (ThermoFisher, 21103049) mixed 1:1 with DMEM/F12 (Gibco, 11320-033), supplemented with 1% B27 (ThermoFisher, 17504044), 1% N-2 (ThermoFisher, 17502048), 1% GlutaMAX™ (Gibco, 35050), 200 µM ascorbic acid (Nacalai Tesque, 03420-52), insulin 7 µg/mL (WAKO, 097-06474), BDNF 10 ng/mL (BioLegend, 788902), and GDNF 10 ng/mL (BioLegend, 760402). The cells were left to form sphere-like aggregates (neurospheres); half of the medium was changed every other day. Neurospheres were removed from the U-bottom wells and attached to the flat surface on culture plastic slides (Ibidi, IB80606) for the differentiation experiment. Neurospheres were also used for the MEA recording and for the in vivo experiment, the results of which are presented in Figure 5.

### Real-time quantitative PCR analysis

RT-qPCR was performed using THUNDERBIRD™ Next SYBR^®^ (TOYOBO, QPX-201) and specifically designed primers (listed in Supplementary Table S1) on QuantStudio™ 3 and QuantStudio™ 7 Flex Real-Time PCR Systems (Applied Biosystems), as described previously ^18^. Data from three biological replicates were analyzed to determine the relative fold change using the 2^−ΔΔCT^ method. Graphs were created using the GraphPad Prism 9 software.

### Immunocytochemistry

Hb-LiNSCs and/or attached neurospheres were fixed with 4% paraformaldehyde and then stained with antibodies against SOX1 (Cell Signaling, 4194S), SOX2 (R&D systems, MAB2018), NESTIN (R&D systems, MAB1259), NANOG (R&D systems, AF1997), POU5F1 (Santa Cruz Biotechnology, sc-5279), PAX6 (Abcam, EPR15858), TUBB3 (GeneTex, GTX85469), OLIG2 (GeneTex, GTX132732), or GFAP (Santa Cruz Biotechnology, sc-33673), and with DAPI for the nuclei. The stained cells were visualized using a Keyence microscope (BZ-X710 or BZ-X810) and an Olympus 3000 confocal microscope system.

### RNA sequencing

For the bulk RNA-seq, cell lysis and RNA extraction and sequencing were performed as described previously ^18^. For analyzing the count data, we used the DESeq2 method for normalization and for differential gene expression analysis^19^. To generate the plots, we used “EnhancedVolcano,” “ComplexHeatmap,” “ggVennDiagram,” “ggplot2,” and other standard RStudio packages. For pathway enrichment, we used Enrichr^20^. As for scRNA-seq, we processed 5000 cells from each sample and prepared the sequencing library using the Chromium Next GEM Single Cell 3′ Reagent Kits v3.1 from 10x Genomics, according to the standard procedures recommended by the manufacturer (Dual Index). After cDNA synthesis, size (TapeStation) and concentration (Qubit) measurements were performed. The size (TapeStation) and concentration (qPCR method) were performed after library preparation. The processing, filtration, analysis, and plotting of data were done in Python using Scanpy^21^.

### Multielectrode array recording

High-density multiple electrode arrays (HD-MEAs) from MaxWell Biosystems were used for detecting neuronal firing and network activity in the neurospheres. One-week-old neurospheres were plated directly onto the iMatrix-511-coated electrode after formation in suspension. Recording was done after six weeks on a MaxOne HD-MEA system. The same electrodes were recorded again after 1 h of 1 µM TTX treatment. The data were analyzed using the MaxLab Live software.

### In vivo neurosphere implantation

Approximately 4-week-old mice were used for the transplantation of one-week-old neurospheres. The neurospheres (about five spheres) were collected with a Hamilton syringe using a stereotactic device and then delivered into the hindbrain region of the mouse brain after drilling the cranium, referring to the coordinates from the mouse brain atlas. After 8 weeks of transplantation, mice were sacrificed according to the ethical guidelines, and then perfusion fixation was performed. Whole brain samples were collected and fixed with 4% paraformaldehyde and then incubated in 30% sucrose for 24 h. The samples were then embedded into a freezing compound for cryosectioning. Thirty-micrometer-thick sections were cut in the sagittal plane of the transplanted hemispheres.

### Statistical analysis

Unpaired parametric Welch’s *t*-test was performed using the GraphPad Prism 9 software. Significance levels were defined as n.s. > 0.05, \**p* ≤ 0.05, \*\**p* ≤ 0.01, \*\*\**p* ≤ 0.001, and \*\*\*\**p* ≤ 0.0001.

