## Supplementary for "Induction and long-term maintenance of hindbrain-like neural stem cells in xeno- and basic fibroblast growth factor-free conditions"

| Gene | forward | reverse |
|---|---|---|
| *CDH1* | AGCAGAACTAACACACGGGG | ACCCACCTCTAAGGCCATCT |
| *CDH2* | CATCCAGACCGACCCAAACA | ACAGACACGGTTGCAGTTGA |
| *FEV* | GACCGAAGCTCCCTCAATCC | CGGGACCTGGAGACCTAGAA |
| *FOXA2* | TGTTCGAGAACGGCTGCTAC | CCCCGAGTTGAGCCTGTGAG |
| *GBX2* | ACTTTTCGCCTCTCGCTGG | AGTCTATGCTGAAGGCGGTG |
| *hACTB* | AGGTCTTTGCGGATGTCCACGT | CACCATTGGCAATGAGCGGTTC |
| *HOXB4* | CCTCGACACCCGCTAACAAA | TATCGGGAGTGGGGGACAAA |
| *HOXB9* | TACCTCACCAGGGACCGTAG | GGGAGGACTGGGGGTAATCT |
| *PAX6* | ATGAGGCTCAAATGCGACTT | GGCCCTTCGATTAGAAAACC |
| *POU5F1* | GCAAAGCAGAAACCCTCGTG | GATCTGCTGCAGTGTGGGT |
| *SIX3* | CTGCCCACCCTCAACTTCTC | CGTGTTTGTTGATGGCCTCG |
| *SLC6A4* | TTGGACGTGTGAGGATGTGG | TCCCTGTTCTCTCCTACGCA |
| *SMA* | TCCGGAGCGCAAATACTCTG | CCCGGCTTCATCGTATTCCT |
| *SOX1* | ATACTGGAGACGAACGCCG | AACCCAAGTCTGGTGTCAGC |
| *SOX2* | GGATAAGTACACGCTGCCCG | ATGTGCGCGTAACTGTCCAT |
| *TPH2* | TATCCCACTCATGCTTGCCG | ACCGTGAAGCCAGACCTTTC |

**Supplementary table 1.**

Primers used in this paper in alphabetical order

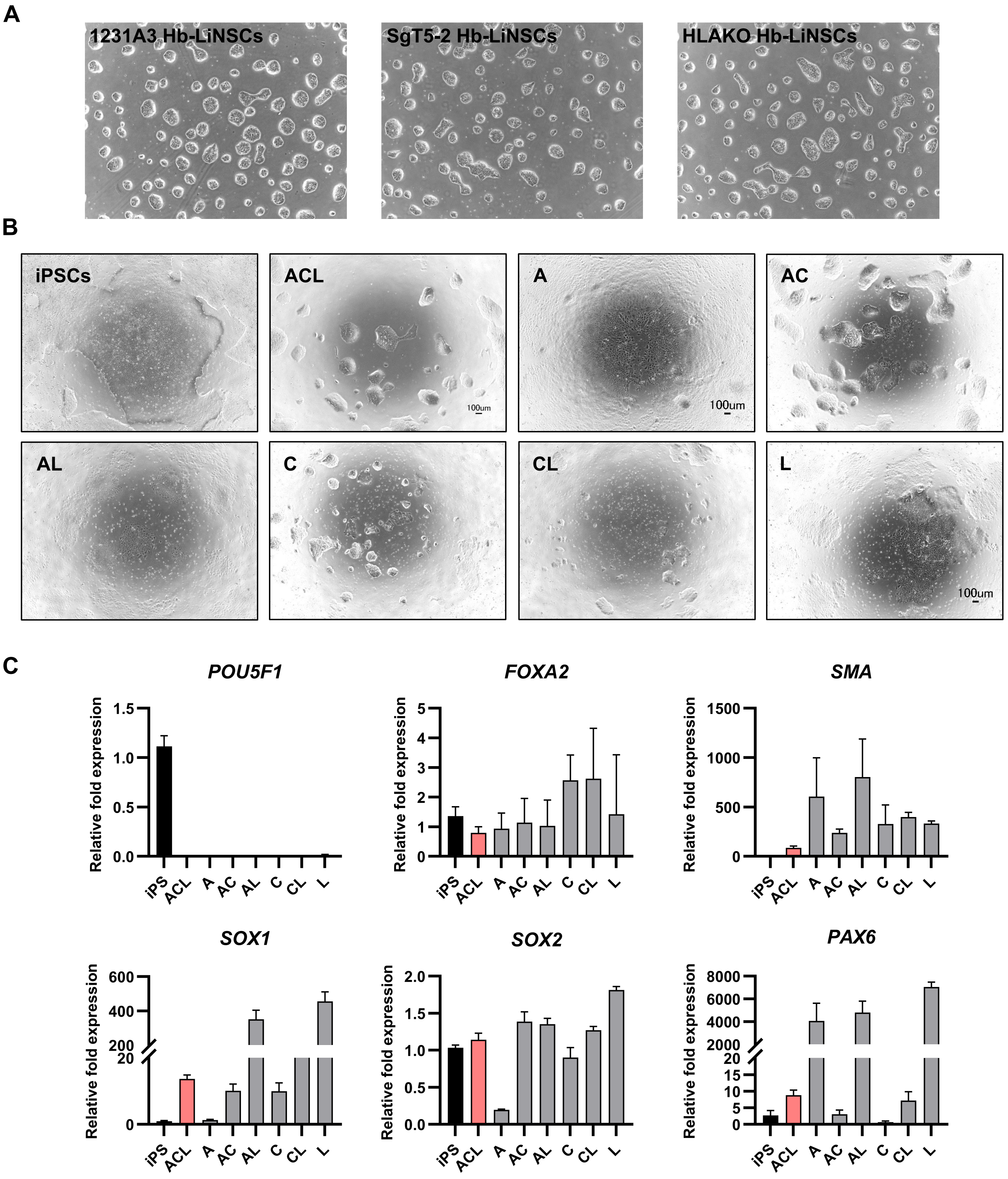


**Supplementary figure 1.**

(A) Phase contrast images of the Hb-LiNSCs induced from three iPSCs lines in the ACL condition at day 7.

(B) Phase contrast images of the iPSCs or iPSCs after one week of induction with A-83-01, CHIR9902, and LDN193189 individually or in combination.

(C) Bar plots of gene expression relative to the iPSCs for the samples in (A) for pluripotency (*POU5F1*), endodermal (*FOXA2*), mesodermal (*SMA*), and ectodermal markers (*SOX1*, *SOX2*, and *PAX6*).

A

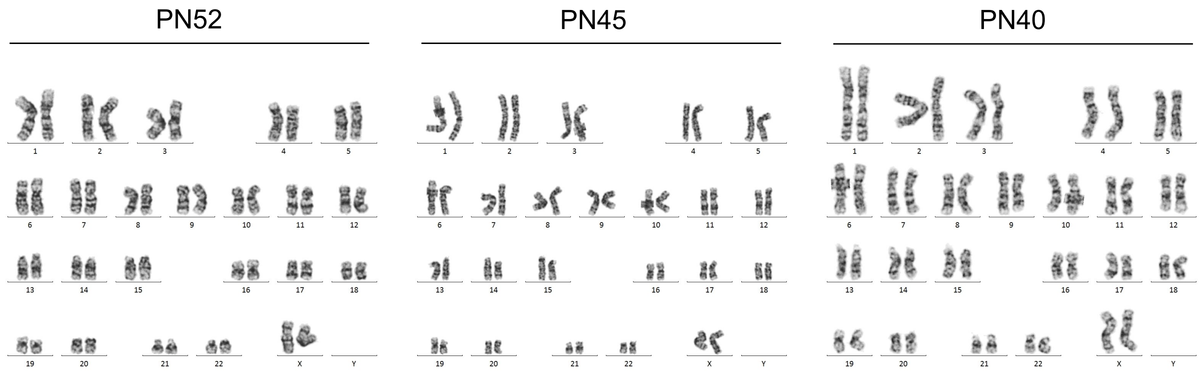


B

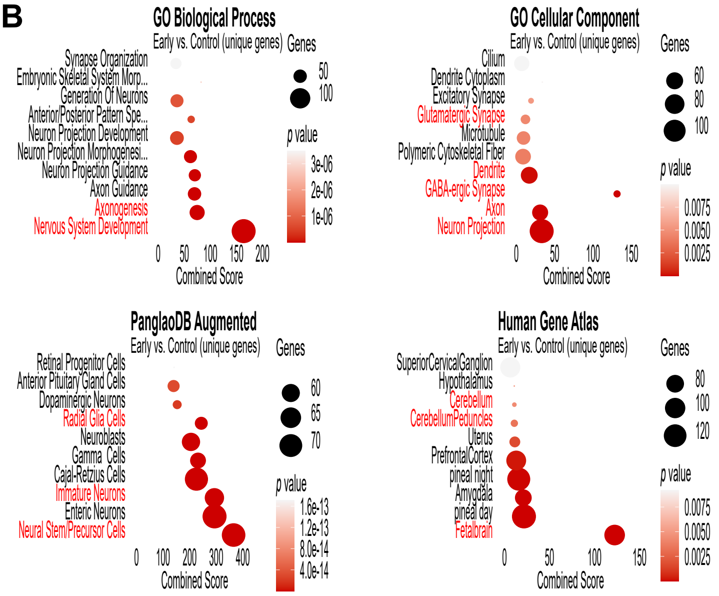


C

| | Total SigDEGs | Upregulated (LFC=>1.5) | Downregulated (LFC=<-1.5) |
|---|---|---|---|
| Early vs. Late | 1225 | 408 | 63 |
| Early vs. Mid | 547 | 207 | 11 |
| Late vs. Mid | 153 | 8 | 45 |
| Early vs. Control | 10467 | 2413 | 2390 |
| Late vs. Control | 10146 | 2206 | 2465 |
| Mid vs. Control | 9886 | 2218 | 2453 |

D

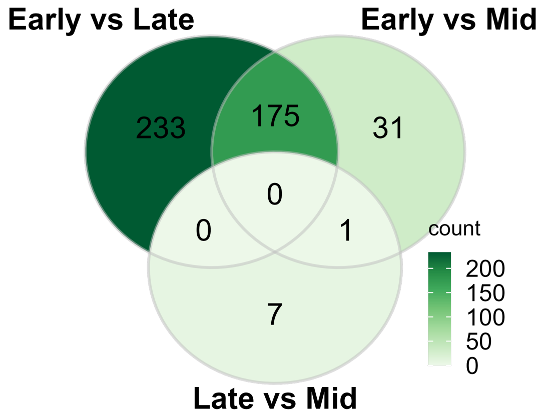


E

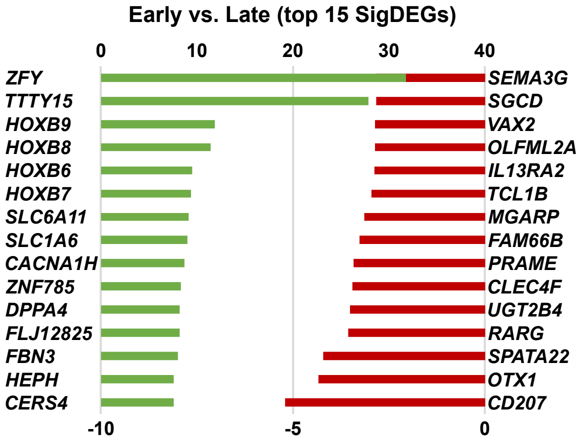


**Supplementary figure 2.**

(A) Hb-LiNSCs karyotyping at PN52, 45, and 40 respectively. Chromosome 9 inversion.

(B) Gene enrichment analysis of the significantly ($p < 0.05$) upregulated (log2foldchange ≥ 1.5) genes in early PN group vs. iPSCs control for selected data sets. Dot size indicates the number of genes overlapped with the data set, color intensity indicates the significance (top 10 terms ordered by p value), and x axis indicates the combined score as calculated by Enrichr.

(C) Number of SigDEGs in the comparative analysis between samples from the bulk RNAseq data.

(D) Venn diagram of the SigDEGs in early vs. mid, late vs. mid, and early vs. late.

(F) Bar plot of the top 15 SigDEGs in early vs. late. The top x-axis is to the left y-axis, and the bottom x-axis is to the right y-xis

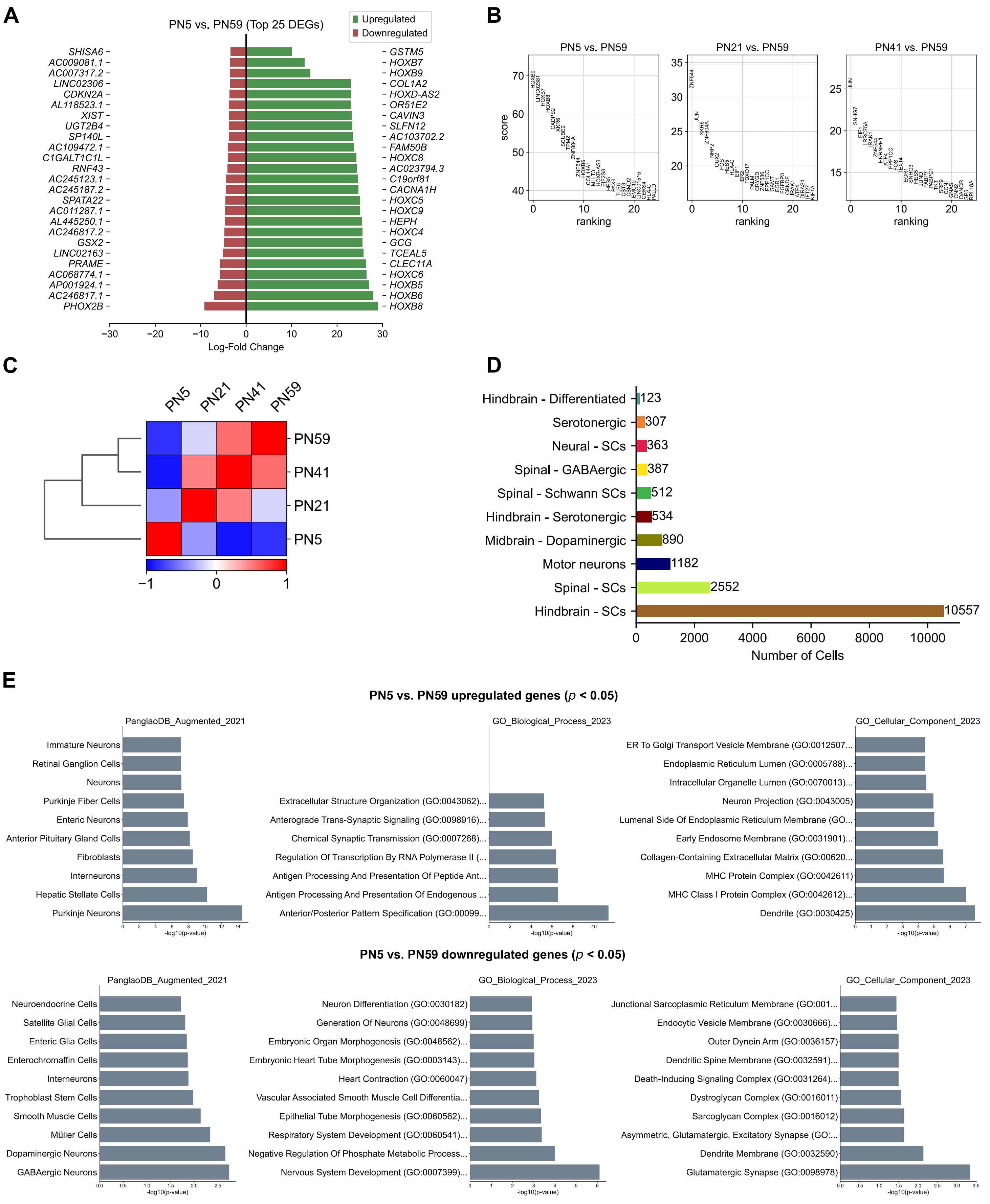


**Supplementary figure 3.**
(A) Bar plot for the top DEGs from the scRNAseq data in PN5 vs PN59 samples
(B) Top genes ranked in the scRNAseq samples when compared to PN59 sample based on the Wilcoxon rank-sum test.
(C) Correlation heatmap between the scRNAseq samples.
(D) Bar plots of the top terms enriched for the indicated terms on top of each plot in the upregulated genes (top panel) and downregulated genes (bottom panel)

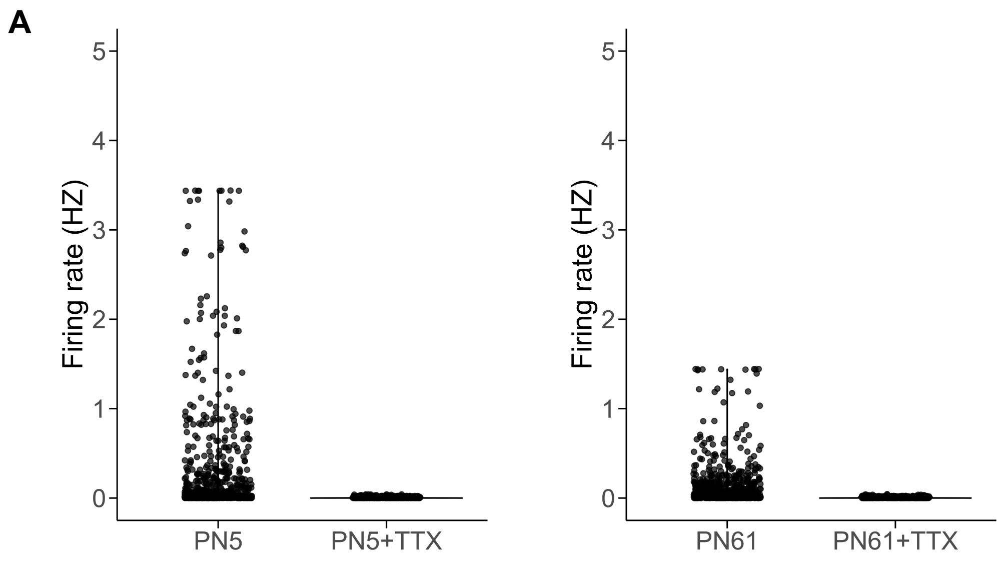


**Supplementary figure 4.**
(A) Firing rate for the recorded electrode in MEA before and after adding TTX. Each dot represents the rate of one of the recorded electrodes.
